# Miffi: Improving the accuracy of CNN-based cryo-EM micrograph filtering with fine-tuning and Fourier space information

**DOI:** 10.1101/2023.12.08.570849

**Authors:** Da Xu, Nozomi Ando

**Affiliations:** Department of Chemistry and Chemical Biology, Cornell University, Ithaca, NY 14850, USA

**Keywords:** cryo-EM, micrograph filtering, CNN, fine-tune, deep learning, software

## Abstract

Efficient and high-accuracy filtering of cryo-electron microscopy (cryo-EM) micrographs is an emerging challenge with the growing speed of data collection and sizes of datasets. Convolutional neural networks (CNNs) are machine learning models that have been proven successful in many computer vision tasks, and have been previously applied to cryo-EM micrograph filtering. In this work, we demonstrate that two strategies, fine-tuning models from pretrained weights and including the power spectrum of micrographs as input, can greatly improve the attainable prediction accuracy of CNN models. The resulting software package, Miffi, is open-source and freely available for public use (https://github.com/ando-lab/miffi).

## Introduction

Single particle cryo-electron microscopy (cryo-EM) is a technique that can provide detailed structural information on the architecture of biological macromolecules with resolution ranging from atomic details to quaternary arrangement (Nogales and Scheres, 2015). It particularly excels at characterizing large biological assemblies or heterogeneous samples, which are extremely challenging targets for other high-resolution structural techniques such as X-ray crystallography or nuclear magnetic resonance (NMR) spectroscopy. Since the introduction of direct electron detection (DED) cameras roughly a decade ago, cryo-EM has undergone rapid development both on the hardware and software fronts (Chua et al., 2022). Improved automation and throughput during data collection, coupled with efficient and user-friendly data processing software, has enabled wide adoption of this technique in many areas of biological research.

Despite the achievements so far, many aspects of cryo-EM can still benefit from further improvements. One such area concerns efficient recognition and exclusion of non-ideal cryo-EM micrographs, given the increasing number and size of cryo-EM datasets. Currently, a beam-image shift scheme (Cheng et al., 2018) is commonly used during cryo-EM data collection using software such as SerialEM (Mastronarde, 2005), EPU, and Leginon (Cheng et al., 2021), which greatly increases data collection throughput. Coupled with newer detectors with shorter exposure times, 300 or more cryo-EM movies can be obtained per hour with little compromise on achievable resolution (Fréchin et al., 2023; Peck et al., 2022). Effort is also being made to automate microscope operation through deep learning methods (Bouvette et al., 2022; Cheng et al., 2023; Fan et al., 2024), which would further accelerate the data collection process. Such advances, combined with the need for a large amount of data for low-concentration or highly heterogeneous samples, have led to increasingly large dataset sizes, typically ranging from a few thousand to tens of thousands of movies. Inevitably, a portion of the collected data will be non-ideal for processing and will thus need to be excluded. A common method for performing filtering is based on contrast transfer function (CTF) fitting, in which the user defines a threshold for acceptable CTF fit resolution or defocus value for a given micrograph. This method is very efficient at excluding micrographs with drift issues or beam aberrations, but it is not ideal for filtering out many other issues such as crystalline ice or off-target support film images. To achieve high filtering accuracy, manual inspection of micrographs is often needed, which is slow and requires expert knowledge to perform. Such an approach becomes less practical as the dataset size increases.

A convolutional neural network (CNN) is a machine learning algorithm that is specialized for processing matrix-like data, such as images. CNNs have been successfully applied to a wide range of computer vision tasks such as image recognition, classification, and segmentation (Alzubaidi et al., 2021). CNNs have also proven useful in many cryo-EM processing steps such as particle picking (Bepler et al., 2019; Tegunov and Cramer, 2019; Wagner et al., 2019), 2D class selection (Kimanius et al., 2021; Li et al., 2020), and micrograph segmentation (Sanchez-Garcia et al., 2020). Because cryo-EM micrograph filtering is an image classification task at its essence, a CNN-based approach is highly attractive. Cianfrocco and co-workers were the first to demonstrate such an approach (Li et al., 2020). By training a CNN based on a ResNet34 model on a labeled dataset of “good” and “bad” micrographs, they were able to show that CNN-based micrograph filtering greatly improves prediction accuracy (~93%) over traditional CTF-based filtering (~78%), in particular by dramatically reducing the false negative rate (“good” micrographs predicted as “bad”). The improvement over CTF-based filtering is especially significant for the classification of tilted images as tilting inherently leads to lower CTF resolution. A more recent version of their tool, MicAssess 1.0 (Li and Cianfrocco, 2021), implements a hierarchical process to further subclassify the “good” and “bad” classes with ~75 and 80% subclass prediction accuracy, respectively. The success of the CNN-based approach to micrograph filtering raises new questions. For example, can we further improve the accuracy of CNN-based filtering given limited training data? Additionally, can we better encode the reasoning behind micrograph filtering into CNN-based methods?

In this work, we examined various strategies to improve CNN-based filtering. First, we tested whether using a CNN pretrained on the ImageNet dataset (Deng et al., 2009), which does not resemble cryo-EM micrographs, as a starting point for training can achieve better micrograph filtering accuracy than training a CNN on cryo-EM data from scratch. This method, known as fine-tuning or transfer learning, has been shown to greatly improve attainable accuracy of CNN models when only limited training data is available (Kolesnikov et al., 2019; Yadav and Jadhav, 2019). Second, we examined whether the direct inclusion of Fourier space information as part of the CNN input could result in a better prediction accuracy, since many issues such as crystalline ice or sample drift can be better spotted in the power spectrum than in the real-space image. Finally, we aimed to expand the versatility of CNN-based filtering in terms of the type of samples and detectors that it can be applied to and provide integration with common processing software packages including RELION (Kimanius et al., 2021) and cryoSPARC (Punjani et al., 2017). We named the resulting tool Miffi, which stands for cryo-EM <u>mi</u>crograph <u>f</u>iltering utilizing <u>F</u>ourier space information (Figure 1). Miffi is open-source and freely available for public use (https://github.com/ando-lab/miffi). Importantly, we find that fine-tuning provides significant improvement over training from scratch and that inclusion of power spectra as a second input channel suppresses the false positive rate (“bad” micrographs predicted as “good”), largely through improved detection of micrographs with crystalline ice. While we provide Miffi for public use, our results also indicate that CNNs pretrained on the ImageNet dataset provide a useful starting point for any user interested in training a model on custom datasets.

**Figure 1.**
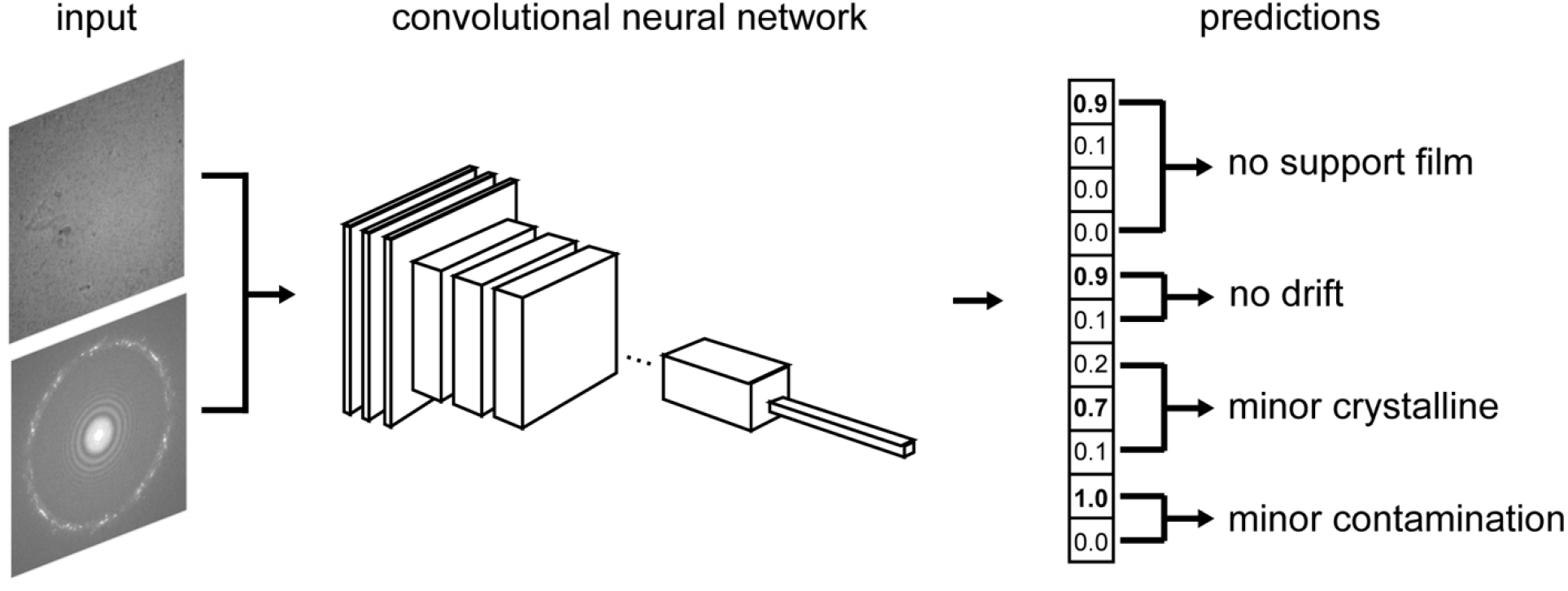
Schematic of Miffi. Miffi employs a convolutional neural network (CNN) for classification by predicting multiple labels for each micrograph. Real-space cryo-EM micrographs and their corresponding power spectra are preprocessed and input into the CNN as two channels, from which an output array containing prediction probability for each label is obtained. Predicted labels are then determined as the one with the highest probability within each of the four label categories: support film coverage, drift, crystallinity, contamination (examples of which can be found in Figure 2).

## Methods

### 1. Data used for training, validation, and testing

All micrographs used in this work were either directly obtained from EMPIAR (Iudin et al., 2023) or obtained in-house and motion corrected using MotionCor2 (Zheng et al., 2017) in RELION 4 (Kimanius et al., 2021). For data collected on a Gatan K3 detector, 7 by 5 patches were used for motion correction, while 5 by 5 patches were used for data collected on a Gatan K2 Summit detector. The training set included a mix of in-house datasets and EMPIAR entries 10202 (Tan et al., 2018), 10249 (Herzik et al., 2019), and 11228 (Filman et al., 2019), as described in Table 1. An in-house dataset on a 240 kDa globular protein collected on a Gatan K3 detector that was not part of the training set was used as the validation set, as it contained a number of micrographs with various issues. All in-house datasets used in training and validation were curated from their respective full sets such that bad micrographs constituted roughly 50% of each set. Micrographs from EMPIAR entries 10175 (Noble et al., 2018b), 10344 (Campbell et al., 2020), 10379 (Li et al., 2020), 10916 (Yang et al., 2022), and 11093 (Röder et al., 2020) were used as additional test sets to evaluate the trained model, the details of which can be found in Table 2. Dose-weighted micrographs were used for all cases.

**Table 1.** Datasets used in the training set. All in-house datasets were curated such that bad micrographs consisted of roughly half of each set. The training set contains micrographs from various types of samples, detectors, and modes of data collection.

| source | particle | notes | detector | num of square micrographs |
| --- | --- | --- | --- | --- |
| in-house | 240 kDa globular protein |  | K3 | 11732 |
| in-house | 300 kDa globular protein | on graphene oxide film, includes tilted images | K3 | 11820 |
| in-house | 210 kDa elongated protein |  | K3 | 3384 |
| in-house | 200 kDa globular protein |  | K3 | 1638 |
| in-house | filament with 160 kDa asymmetric unit |  | K3 | 3712 |
| in-house | methionine synthase mixed with apoferritin (Watkins et al., 2023) | includes tilted images | K3 | 10050 |
| EMPIAR-10202 | adeno-associated virus (Tan et al., 2018) | 3.8 MDa globular | K2 Summit | 1317 |
| EMPIAR-10249 | alcohol dehydrogenase (Herzik et al., 2019) | 81 kDa globular, very densely packed | K2 Summit | 1151 |
| EMPIAR-11228 | cytochrome OmcS nanowire (Filman et al., 2019) | thin filament with 50 kDa asymmetric unit | K2 Summit | 964 |

**Table 2.** Additional test sets. The test sets contain a diverse set of micrographs that were not part of the training set.

| source | particle | notes | detector |
| --- | --- | --- | --- |
| EMPIAR-10175 | hemagglutinin (Noble et al., 2018b) | 190 kDa elongated | K2 Summit |
| EMPIAR-10344 | αVβ8 integrin (Campbell et al., 2020) | 240 kDa elongated on graphene oxide film | K2 Summit |
| EMPIAR-10379 | aldolase (Li et al., 2020) | 160 kDa globular | K2 Summit |
| EMPIAR-10916 | amyloid-β 42 filament (Yang et al., 2022) | filament with 9 kDa asymmetric unit | Falcon 4i |
| EMPIAR-11093 | amylin amyloid fibril (Röder et al., 2020) | thin filament with 8 kDa asymmetric unit | Falcon III |

To maximize applicability of the model on data collected from different detectors which are of different dimensions, we set the CNN inputs as square images. Micrographs that are non-square were thus split or cropped into square micrographs during the preprocessing stage for training and inference. For analyses done in this work, micrographs from a Gatan K3 detector (5760 pix × 4092 pix) were split into two square micrographs (4092 pix × 4092 pix) spanning the original micrograph with overlap in the middle (2424 pix × 4092 pix). Micrographs from a Gatan K2 Summit detector, which are nearly square (3838 pix × 3710 pix), were cropped into a single square micrograph (3710 pix × 3710 pix) starting from the left edge of the original micrograph oriented with the long dimension in the horizontal direction. Micrographs that are originally square (e.g., collected on a TFS Falcon detector) were kept as is. Each square micrograph in the training and validation sets was labeled individually and treated as separate entries. The training set included a total of 45768 square micrographs, while the validation set included a total of 4000 square micrographs.

### 2. Labeling scheme

The training, validation, and test datasets were labeled manually by inspecting each square micrograph along with its corresponding power spectrum. A GUI interface designed for this process is included in the GitHub repository. Each square micrograph was given four labels representing different categories of issues: support film coverage, drift, crystallinity, contamination (Figure 1). The first label describes the degree of the support film coverage in the micrograph, which can be one of the following: no film, minor film, major film, film (Figure 2, top row). Note that film here only means support film of the grid itself, and additional continuous film such as mono-layer graphene oxide is not considered film in this case. The second label is binary and describes issues of sample displacement, where sample drift, cracks, or an empty micrograph is labeled as “bad” (Figure 2, second row). The third label describes the crystallinity of the ice in the sample, which is determined by non-diffuse intensity at around 1/3.7 Å^-1^ in Fourier space, and can be one of the following: not crystalline, minor crystalline, major crystalline (Figure 2, third row). Finally, the fourth label is binary and describes whether the micrograph is covered largely in contaminant objects, which includes ice crystals and ethane contamination (Figure 2, bottom row). It is worth noting that as the changes within each category are often continuous, it is difficult to set a clear cutoff, but extra care was made to keep the labeling scheme as consistent as possible.

**Figure 2.**
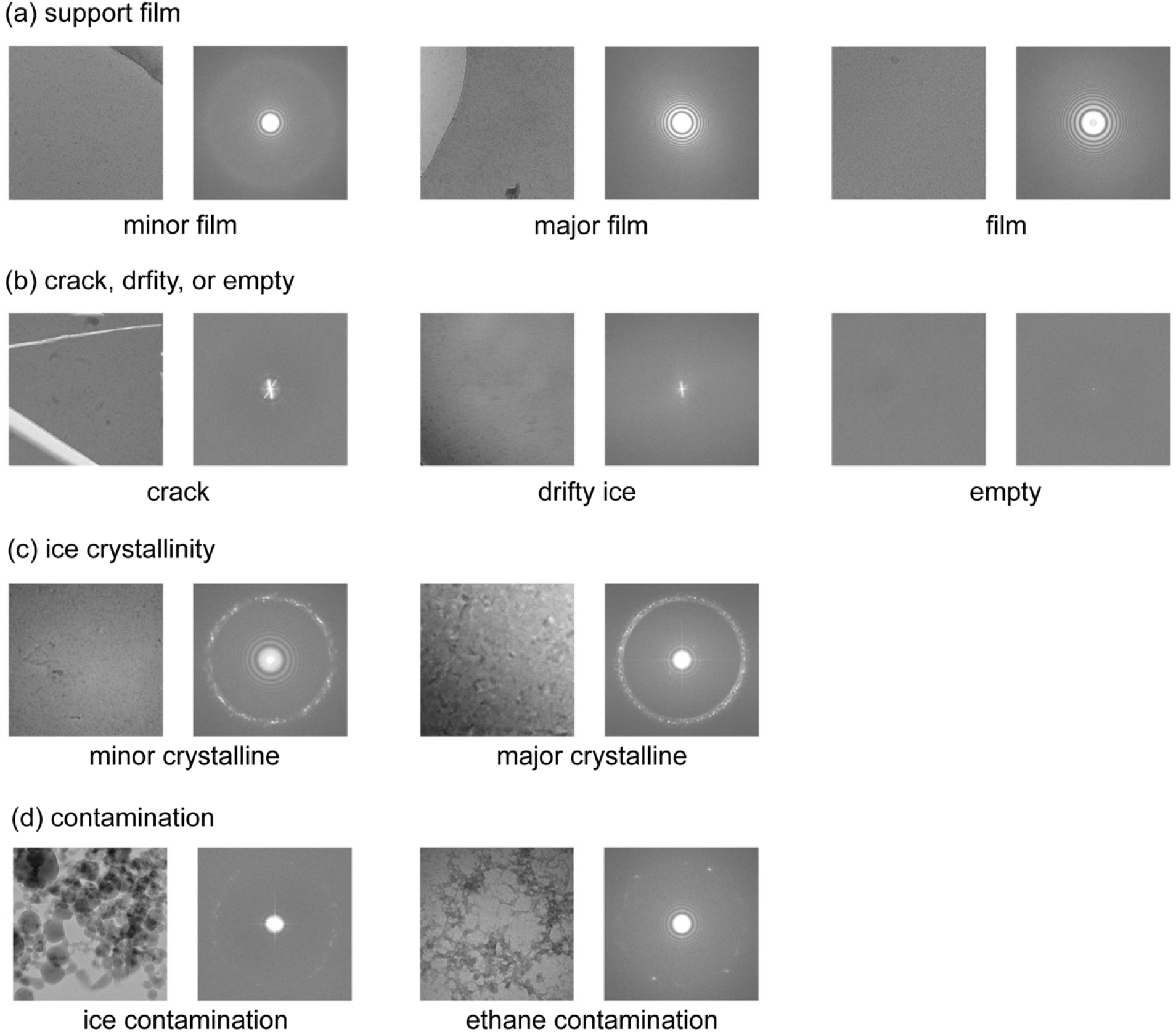
Example “bad” micrographs for each label category from the in-house training set. The real-space micrograph is shown on the left, with its corresponding power spectrum shown on the right. (a) Support film category. Support carbon film in the example is the darker region on the right side of the real-space micrograph, while the ice-containing hole is the lighter region on the left side. (b) Sample displacement category. Translational motion of the sample caused by drift or cracks often results in asymmetric resolution or a streaky pattern in the power spectrum. An empty hole on the other hand has a flat power spectrum with high intensity at the origin of Fourier space. (c) Ice crystallinity category. Ice crystallinity is determined by a non-diffuse ring at 1/3.7 Å^-1^ in Fourier space. Micrographs with strong ice crystallinity often exhibit clear dark-and-light alternating patterns in real space (example on the right). However, for samples with minor crystallinity, features in the real-space micrographs are often subdued and thus are better identified with the ring feature in the power spectra. (e) Contamination category. Two examples shown here represent contaminating ice crystals adhering to the hole (left) and ethane contaminants embedded in ice (right).

### 3. Model training

The ConvNeXt-Small model (Liu et al., 2022) was chosen for the classification task in this work based on its simplicity in architecture and high performance on the ImageNet datasets. All models and their training were implemented in Pytorch (Paszke et al., 2019). Training was performed either from scratch using randomly initialized model weights, or via fine-tuning of model weights pretrained on the ImageNet-12k dataset (Deng et al., 2009; Liu et al., 2022). We tested both a single-channel input of real-space micrographs as well as a two-channel input of the real-space micrographs with their corresponding power spectra. The Timm package (Wightman, 2019) was used to modify the input and output layers of the model to match the desired number of input channels and output classes. As the ImageNet-pretrained model was trained with three input channels corresponding to three colors, pretrained weights were reduced to one or two channels in the following manner to maintain normalization: for a single input channel, the sum of weights from the original three channels was used; for two input channels, the weights in first two original channels were multiplied by a factor of 1.5, while the third was unused.

Square micrographs were Fourier cropped to 1.5 Å pixel size during preprocessing to keep the location of the ice ring consistent in the power spectrum. During the training stage, data augmentation was subsequently applied, including a random crop with a ratio randomly chosen from 0.8 – 1.0, along with a random horizonal and vertical flip. Data augmentation was not applied during validation or inference. For two-channel input models, the power spectra of the augmented micrographs were computed and appended as an additional channel. Inputs were then downsampled to 384 pix × 384 pix with bilinear interpolation, and finally normalized to a mean of 0 with a standard deviation of 1 for each channel individually, with pixels that have intensity falling outside 2.5 standard deviation thresholded (to ensure that micrographs from different sources are on a similar intensity scale). During training, the four label categories for each micrograph are treated as independent from each other, while the probability for all possible labels within each label category is summed to one using the softmax function. Cross-entropy between true label and predicted label probability (softmax of CNN output) for all four label categories were calculated individually, and the sum of which was defined as the loss function. The AdamW optimizer (Loshchilov and Hutter, 2017) and a cosine learning rate scheduler were used for training all models. Layer-wise learning rate decay was applied in fine-tuning by dividing layers into 12 groups and setting the learning rate for each group. Linear warmup epochs were used in the case of training from scratch. Detailed parameters used in training can be found in Table 3. Notably, the fine-tuning process was found to be prone to overfitting, likely reflecting the fact that our training set is small relative to the size of the CNN such that the models can overfit and lose generality. The learning rate, number of training epochs, and layer decay ratio were thus chosen carefully to minimize overfitting, which can be visualized in the loss functions (i.e., the training loss will continue to decrease as the model improves its fit with the training set, but the validation loss will begin to increase as the model begins to lose its generality). All models were trained on a NVIDIA RTX-3090 GPU with less than an hour per epoch.

**Table 3.** Hyperparameters used in model training.

|  | from scratch<br>single-channel | fine-tune<br>single-channel | fine-tune<br>two-channel |
| --- | --- | --- | --- |
| optimizer | AdamW | AdamW | AdamW |
| base learning rate | 4e-4 | 1e-5 | 1e-5 |
| weight decay | 0.05 | 1e-8 | 1e-8 |
| batch size | 32 | 32 | 32 |
| training epochs | 20 | 9 | 10 |
| warmup epochs | 5 | None | None |
| learning rate (lr) schedule | cosine decay | cosine decay | cosine decay |
| layer lr decay ratio | None | 0.9 | 0.9 |

To test how the number of training samples can affect achievable accuracy, trainings were performed with subsets of the full training set in the following manner. The full training set was downsampled to 30000, 20000, 10000, 5000, 2000, 1000, 500, 200, 100 micrographs, by randomly removing samples while keeping the relative number of different labels the same to ensure diversity in training samples. Two-channel ConvNeXt-Small and ConvNeXt-Tiny models pretrained on ImageNet data were then fine-tuned on the downsampled training sets with the same hyperparameters as the two-channel fine-tuned model in Table 3 for 10 epochs, with intermediate models saved at each epoch. The intermediate model with the highest validation accuracy in each training was then used to perform the inference on the validation set to calculate the good-versus-bad prediction accuracy.

### 4. Inference and calculation of prediction accuracy

During inference, micrographs were first split into square micrographs and Fourier cropped to 1.5 Å pixel size as described above for the training datasets. Power spectra were then appended when applicable, followed by downsampling and normalization, as were done in training. The prepared input was passed through the CNN model to obtain the output array, which was then converted to probabilities for each label using the softmax function. The predicted label for each category was determined as the one with highest probability (Figure 1, right). For micrographs with multiple square splits, predictions from individual splits are combined with a predefined set of rules which can be customized by the user (e.g., the user can choose to only keep micrographs in which all splits are classified as “good,” or a user may choose the maximize the number of micrographs by also keeping ones that are partially “bad”). In this work, because each square split was labeled individually, accuracy calculation was also performed for each split without combining. For the good-versus-bad accuracy calculations shown here, we defined the following for the two non-binary categories: for the support film coverage category, we defined “good” micrographs as ones that are labeled as either no film or minor film; for the crystalline ice category, we defined “good” micrographs as ones that are labeled as not crystalline. Overall “good” micrographs were then defined as ones that are labeled as “good” in all four categories, with the rest defined as bad micrographs. Accuracy was calculated as the number of correct predictions divide by the total number of samples. True positive denotes the number of correctly predicted “good” micrographs, while true negative denotes the number of correctly predicted “bad” micrographs. False positive denotes the number of “bad” micrographs predicted as “good”, while false negative denotes the number of “good” micrographs predicted as “bad”.

To compare the performance of our model with existing methods, micrograph filtering was performed with MicAssess (Li et al., 2020; Li and Cianfrocco, 2021) and CTF-based filtering using the same test sets. Version 1.0 of MicAssess was obtained from GitHub, and the model weights were obtained from the authors. Default parameters were used during inference, and micrographs predicted as “good” in the binary classification step of MicAssess were treated as “good” micrographs, while the rest were treated as “bad” micrographs. To compare our model with CTF-based filtering, patch CTF estimation was performed in cryoSPARC (Punjani et al., 2017). Because the distribution of CTF fitting resolution varied greatly between different datasets, we set the cutoff individually for each dataset. Micrographs with CTF fitting resolution worse than 10 Å were first excluded as they greatly biased the statistics and were treated as “bad” micrographs. Of the remainder, those with CTF fitting resolution within two standard deviations of the mean were treated as “good” micrographs, while those outside of this range were treated as “bad” micrographs.

## Results and Discussions

We tested two strategies in model design and training and evaluated their effects on the prediction accuracy with the validation set. The first strategy was to perform fine-tuning starting from model weights pretrained on the ImageNet-12k dataset, which contrasts with previous work which trained a model from scratch (Li et al., 2020). Although the ImageNet dataset consists of everyday objects that are very different from cryo-EM images, previous examples such as medical imaging have shown that pretrained weights can still notably improve accuracy especially in cases where limited training data is available (Yadav and Jadhav, 2019). The second strategy was to directly include power spectra of the micrographs as an additional channel in the input. This is inspired by the fact that many issues with micrographs can be spotted more easily in the power spectrum, particularly the ice crystallinity. Although information in the Fourier space representation should theoretically also be present in the real-space images, the information content will be significantly reduced after the downsampling process, which is a necessary preprocessing step when utilizing CNNs on micrographs due to computational cost. By explicitly including power spectra as part of the input, high-resolution Fourier space information contained in the original micrographs can be preserved through the downsampling step. Here, we compared the resulting models of three training cases: (1) training from scratch with a single-channel real-space input, (2) fine-tuning from a pretrained model with a single-channel real-space input, and (3) fine-tuning with a two-channel input that includes power spectra. The resulting overall good-versus-bad micrograph accuracy can be found in Table 4, while the accuracy for individual categories can be found in Supplementary Table S1.

**Table 4.** Overall accuracy (acc.) of good-versus-bad predictions made for the validation set by the three models. True positive = TP; false positive = FP; true negative = TN; false negative = FN.

|  | acc. | TP | FP | TN | FN |
| --- | --- | --- | --- | --- | --- |
| from scratch<br>single-channel | 0.780 | 1749 | 418 | 1371 | 462 |
| fine-tune<br>single-channel | 0.961 | 2178 | 122 | 1667 | 33 |
| fine-tune<br>two-channel | 0.986 | 2183 | 27 | 1762 | 28 |

By comparing the resulting accuracy from the first two training cases (Table 4, top two rows), we can see that fine-tuning from pretrained weights greatly improved the attainable accuracy with the same training data, from 78% to 96% or greater. This can be rationalized as the feature extracting capability of the initial layers in the CNN being largely transferable even with significant differences between pretraining ImageNet data and cryo-EM images. This is consistent with the observation that layer decay in learning rate improved attainable accuracy (results not shown), indicating that weights in initial layers require less change to arrive at the optimal values. Furthermore, we found that when fine-tuning is performed, the training loss is already low after a single epoch of training, and becomes lower than the final loss of training from scratch after an additional epoch (Figure 3), again suggesting that pretrained weights are much closer to the optimal values compared to randomly initialized weights. The second strategy, direct inclusion of power spectra, additionally increased the prediction accuracy on top of fine-tuning from 96% to 99%, evident from comparison between the last two training cases (Table 4, bottom two rows). When comparing prediction accuracy in individual categories (Supplementary Table S1), the most significant improvement comes from the ice crystallinity category. Notably, the number of false positives in the prediction of ice crystallinity was greatly reduced, consistent with the intuition that power spectra provide better indication for the presence of crystalline ice than real-space images.

**Figure 3.**
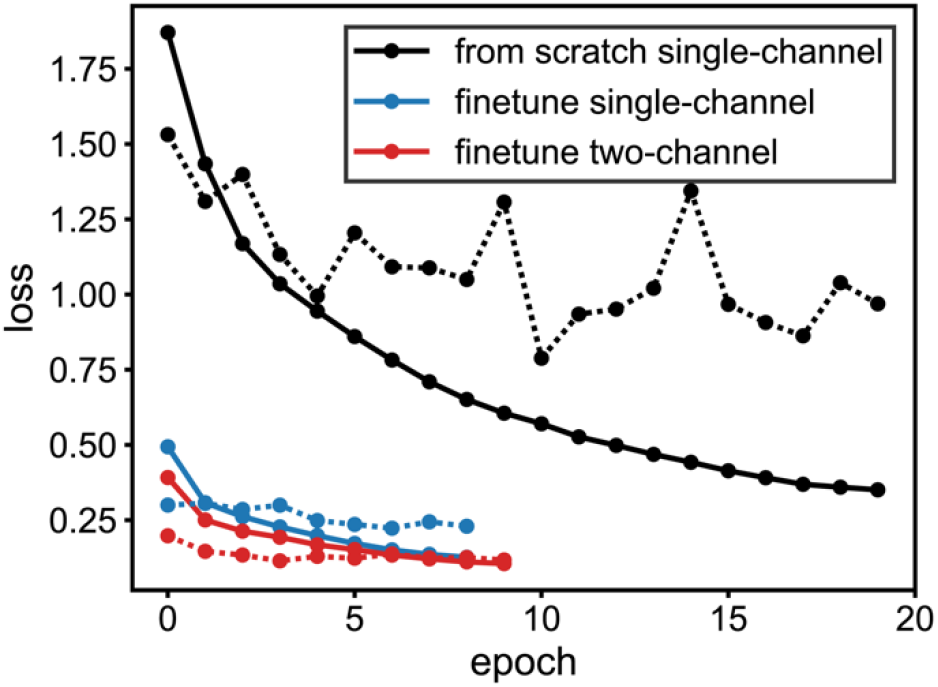
Loss versus epoch for all three training cases. Solid lines denote training loss while dashed lines denote validation loss. Black, blue, and red lines correspond to training from scratch with single-channel input, fine-tuning with single-channel input, and fine-tuning with two-channel input, respectively.

To further examine the generality of our two-channel input model trained with fine-tuning, we tested its performance on five additional datasets from EMPIAR, which contains a variety of features including filamentous samples or graphene oxide films. The resulting prediction accuracies (Table 5 and Supplementary Table S2) show that our model performs relatively well for all chosen datasets, with an overall accuracy higher than 95% in all cases. To compare with other existing methods, we performed micrograph filtering with MicAssess and CTF-fitting on the same EMPIAR datasets and calculated prediction accuracies based on our labels (Supplementary Table S3 and Supplementary Table S4). The results indicate that our model provides the highest filtering accuracy among tested methods for all datasets. Surprisingly, we found that CTF-based filtering outperformed MicAssess for many of the datasets tested here. This could be a result of these test sets differing significantly from the training set for MicAssess. It is also important to note that our labeling criteria may differ from those used in training MicAssess. In the absence of a standardized and validated test set for micrograph filtering, an exact comparison cannot be made between different methods. Nonetheless, these results do illustrate the utility of utilizing Fourier information. A notable example was EMPIAR-10379, for which CTF-based filtering had an overall accuracy of 97.8%, compared to MicAssess, which had 77.5% accuracy. Interestingly, this dataset had very few issues in the ice crystallinity, film, and contamination categories (17 out of 1118 micrographs) but had many in the drift category (218 out of 1118) (Supplementary Table S2), indicating that CTF-based filtering is better at detecting the latter type of issue. This observation is consistent with the fact that CTF fitting is done in Fourier space, and drift issues are readily detectable as they lead to anisotropic power spectra or loss of Thon rings, in the case of empty holes. The fact that our model outperforms both CTF-based filtering and MicAssess here (overall accuracy of 99.6%) further demonstrates that combining information from power spectra and real space micrographs is key to accurately assessing micrograph quality. Future benchmarking studies would benefit from the development of publicly available standardized test set.

**Table 5.** Overall good-versus-bad accuracy (acc.) on additional test sets. True positive = TP; false positive = FP; true negative = TN; false negative = FN.

|  | acc. | TP | FP | TN | FN |
| --- | --- | --- | --- | --- | --- |
| EMPIAR-10175 | 0.954 | 802 | 31 | 171 | 16 |
| EMPIAR-10344 | 0.979 | 2348 | 26 | 278 | 30 |
| EMPIAR-10379 | 0.996 | 884 | 2 | 230 | 2 |
| EMPIAR-10916 | 0.971 | 1684 | 0 | 27 | 51 |
| EMPIAR-11093 | 0.973 | 1123 | 16 | 171 | 20 |

While our model appears to be generally applicable, we do observe slight differences in accuracy for different test sets. We hypothesized that these variations correspond to their degree of dissimilarity to the training set, and attempts were made to improve our model by training it with test sets included as part of the training set and with the same hyperparameters as before. The resulting models, however, show slight deterioration in performance on additional hold-out test sets (results not shown). This could either be due to the new training set requiring additional optimization of hyperparameters, or that training specialized models on individual subsets may be necessary to improve the accuracy beyond what we see here. To investigate how many training samples are required to obtain a reasonable prediction accuracy, we performed training on random subsets of the full training set with various sizes. We also performed the training with both ConvNeXt-Small and ConvNeXt-Tiny models to explore whether a smaller CNN provides better training results when the number of training samples is low. The result, shown in Figure 4, suggests that a relatively high accuracy (~94%) can be achieved with a relatively small training set of only 1000 micrographs, and the accuracy plateaus when the number of training samples is higher than 10000 micrographs. Interestingly, we found that ConvNeXt-Tiny did not appear to provide an advantage over ConvNeXt-Small in the low training sample regime, indicating that ConvNeXt-Small is a suitable choice even when training with very limited data. Overall, our result here suggests that with the fine-tuning strategy, training specialized models should not be difficult to achieve even with a small amount of data.

**Figure 4.**
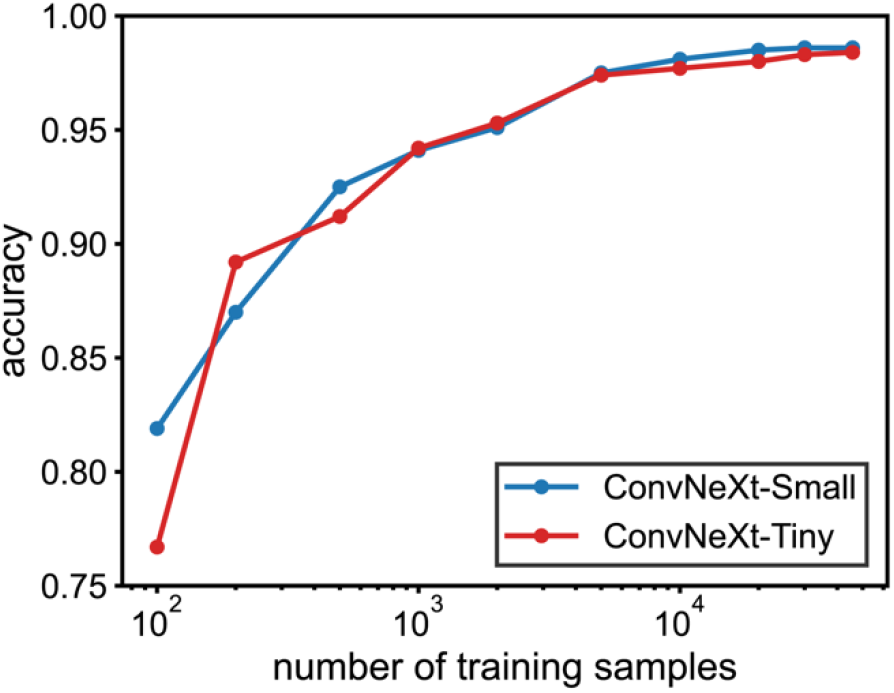
Overall good-versus-bad accuracy on validation set plotted against the number of training samples used in training. Blue and red lines correspond to results from training ConvNeXt-Small and ConvNeXt-Tiny models, respectively.

## Conclusion

In this work, we examined various strategies to improve CNN-based cryo-EM micrograph filtering. We showed that two strategies, fine-tuning from pretrained weights and directly including power spectra as input, can improve the attainable accuracy of resulting models. This likely results from the transferability of the feature-extracting capability in the initial layers of a pretrained model, as well as the existence of better indicators for certain micrograph features in Fourier space. We demonstrated that a model trained with these strategies can filter a diverse set of cryo-EM datasets with high accuracy.

The resulting software, named Miffi, is open-source and freely available for public use. Miffi is implemented with Pytorch, which enables cross-platform compatibility. Miffi can be accelerated with CUDA on a NVIDIA GPU (about 3 micrographs per second with our setup), but it also runs with reasonable speed when using CPU only (about 3 seconds per micrograph with our setup). Miffi accepts micrograph inputs in file formats produced by common cryo-EM processing software, such as RELION and cryoSPARC, and can write output files in the same format such that they can be directly imported back into the originating software. Necessary preprocessing steps (e.g., splitting/cropping micrographs into square micrographs, Fourier cropping, calculation of power spectra, downsampling) are performed on input micrographs in Miffi before passing them to the CNN for classification. Miffi also provides users the flexibility to control the classification process, including the ability to customize rules for combining predictions for micrographs with multiple square splits, to filter predictions based on their confidence scores, and to control which categories are written out. We note that the labeled categories in our training process do not include all potential issues, such as low particle visibility or thick ice. In particular, ice thickness is a continuum quantity that is best measured experimentally (Neselu et al., 2023; Noble et al., 2018a; Rice et al., 2018), for which the ideal range is sample dependent. Therefore, combining Miffi with other data assessment criteria will still be beneficial. For example, issues with ice thickness can be filtered based on experimentally measured values or based on the intensity of the diffuse ice ring at 1/3.7 Å^-1^ in Fourier space, and issues with particle visibility can be filtered by excluding micrographs with low defocus values. Overall, we believe that Miffi can be readily incorporated into common data processing pipelines and greatly improve the accuracy and efficiency of the micrograph curation step.

## Acknowledgements

We thank Dr. Steve Meisburger for critical reading of the manuscript and the following Ando Lab members for collecting data used for training: Dr. Audrey Burnim, Dr. Amanda Byer, Dr. Gabrielle Illava, Rob Miller, Haoyue Wang, Dr. Max Watkins. In-house data used in this work were collected at the Cornell Center for Materials Research (CCMR) Shared Facilities, which are supported through the National Science Foundation (NSF) MRSEC program (DMR-1719875), and the National Center for CryoEM Access and Training (NCCAT) and the Simons Electron Microscopy Center located at the New York Structural Biology Center, supported by the National Institutes of Health (NIH) Common Fund Transformative High Resolution Cryo-Electron Microscopy program (U24 GM129539) and by grants from the Simons Foundation (SF349247) and NY State Assembly. This work was supported by NSF grant MCB-1942668 (to N.A.), NIH grant GM124847 (to N.A.), and startup funds from Cornell University (to N. A.).

## Data and Code Availability

The Miffi package is available on GitHub (https://github.com/ando-lab/miffi). Model weights trained in this work are available on Zenodo (https://doi.org/10.5281/zenodo.8342009). Labels for EMPIAR test sets are available on GitHub.

## References

Alzubaidi, L., Zhang, J., Humaidi, A.J., Al-Dujaili, A., Duan, Y., Al-Shamma, O., Santamaría, J., Fadhel, M.A., Al-Amidie, M., Farhan, L., 2021. Review of deep learning: concepts, CNN architectures, challenges, applications, future directions. J Big Data 8, 53.

Bepler, T., Morin, A., Rapp, M., Brasch, J., Shapiro, L., Noble, A.J., Berger, B., 2019. Positive-unlabeled convolutional neural networks for particle picking in cryo-electron micrographs. Nat. Methods 16, 1153–1160.

Bouvette, J., Huang, Q., Riccio, A.A., Copeland, W.C., Bartesaghi, A., Borgnia, M.J., 2022. Automated systematic evaluation of cryo-EM specimens with SmartScope. Elife 11. 10.7554/eLife.80047

Campbell, M.G., Cormier, A., Ito, S., Seed, R.I., Bondesson, A.J., Lou, J., Marks, J.D., Baron, J.L., Cheng, Y., Nishimura, S.L., 2020. Cryo-EM Reveals Integrin-Mediated TGF-β Activation without Release from Latent TGF-β. Cell 180, 490–501.e16.

Cheng, A., Eng, E.T., Alink, L., Rice, W.J., Jordan, K.D., Kim, L.Y., Potter, C.S., Carragher, B., 2018. High resolution single particle cryo-electron microscopy using beam-image shift. J. Struct. Biol. 204, 270–275.

Cheng, A., Kim, P.T., Kuang, H., Mendez, J.H., Chua, E.Y.D., Maruthi, K., Wei, H., Sawh, A., Aragon, M.F., Serbynovskyi, V., Neselu, K., Eng, E.T., Potter, C.S., Carragher, B., Bepler, T., Noble, A.J., 2023. Fully automated multi-grid cryoEM screening using Smart Leginon. IUCrJ 10, 77–89.

Cheng, A., Negro, C., Bruhn, J.F., Rice, W.J., Dallakyan, S., Eng, E.T., Waterman, D.G., Potter, C.S., Carragher, B., 2021. Leginon: New features and applications. Protein Sci. 30, 136–150.

Chua, E.Y.D., Mendez, J.H., Rapp, M., Ilca, S.L., Tan, Y.Z., Maruthi, K., Kuang, H., Zimanyi, C.M., Cheng, A., Eng, E.T., Noble, A.J., Potter, C.S., Carragher, B., 2022. Better, Faster, Cheaper: Recent Advances in Cryo-Electron Microscopy. Annu. Rev. Biochem. 91, 1–32.

Deng, J., Dong, W., Socher, R., Li, L.-J., Li, K., Fei-Fei, L., 2009. Imagenet: A large-scale hierarchical image database, in: 2009 IEEE Conference on Computer Vision and Pattern Recognition. Ieee, pp. 248–255.

Fan, Q., Li, Y., Yao, Y., Cohn, J., Liu, S., Xu, Z., Vos, S., Cianfrocco, M., 2024. CryoRL: Reinforcement Learning Enables Efficient Cryo-EM Data Collection, in: Proceedings of the IEEE/CVF Winter Conference on Applications of Computer Vision. pp. 7892–7902.

Filman, D.J., Marino, S.F., Ward, J.E., Yang, L., Mester, Z., Bullitt, E., Lovley, D.R., Strauss, M., 2019. Cryo-EM reveals the structural basis of long-range electron transport in a cytochrome-based bacterial nanowire. Commun Biol 2, 219.

Fréchin, L., Holvec, S., von Loeffelholz, O., Hazemann, I., Klaholz, B.P., 2023. High-resolution cryo-EM performance comparison of two latest-generation cryo electron microscopes on the human ribosome. J. Struct. Biol. 215, 107905.

Herzik, M.A., Jr, Wu, M., Lander, G.C., 2019. High-resolution structure determination of sub-100 kDa complexes using conventional cryo-EM. Nat. Commun. 10, 1032.

Iudin, A., Korir, P.K., Somasundharam, S., Weyand, S., Cattavitello, C., Fonseca, N., Salih, O., Kleywegt, G.J., Patwardhan, A., 2023. EMPIAR: the Electron Microscopy Public Image Archive. Nucleic Acids Res. 51, D1503–D1511.

Kimanius, D., Dong, L., Sharov, G., Nakane, T., Scheres, S.H.W., 2021. New tools for automated cryo-EM single-particle analysis in RELION-4.0. Biochem. J 478, 4169–4185.

Kolesnikov, A., Beyer, L., Zhai, X., Puigcerver, J., Yung, J., Gelly, S., Houlsby, N., 2019. Big Transfer (BiT): General Visual Representation Learning. arXiv [cs.CV].

Li, Y., Cash, J.N., Tesmer, J.J.G., Cianfrocco, M.A., 2020. High-Throughput Cryo-EM Enabled by User-Free Preprocessing Routines. Structure 28, 858–869.e3.

Li, Y., Cianfrocco, M., 2021. MicAssess https://github.com/cianfrocco-lab/automatic-cryoem-preprocessing.

Liu, Z., Mao, H., Wu, C.-Y., Feichtenhofer, C., Darrell, T., Xie, S., 2022. A ConvNet for the 2020s. arXiv [cs.CV].

Loshchilov, I., Hutter, F., 2017. Decoupled Weight Decay Regularization. arXiv [cs.LG].

Mastronarde, D.N., 2005. Automated electron microscope tomography using robust prediction of specimen movements. J. Struct. Biol. 152, 36–51.

Neselu, K., Wang, B., Rice, W.J., Potter, C.S., Carragher, B., Chua, E.Y.D., 2023. Measuring the effects of ice thickness on resolution in single particle cryo-EM. J Struct Biol X 7, 100085.

Noble, A.J., Dandey, V.P., Wei, H., Brasch, J., Chase, J., Acharya, P., Tan, Y.Z., Zhang, Z., Kim, L.Y., Scapin, G., Rapp, M., Eng, E.T., Rice, W.J., Cheng, A., Negro, C.J., Shapiro, L., Kwong, P.D., Jeruzalmi, D., des Georges, A., Potter, C.S., Carragher, B., 2018a. Routine single particle CryoEM sample and grid characterization by tomography. Elife 7. 10.7554/eLife.34257

Noble, A.J., Wei, H., Dandey, V.P., Zhang, Z., Tan, Y.Z., Potter, C.S., Carragher, B., 2018b. Reducing effects of particle adsorption to the air-water interface in cryo-EM. Nat. Methods 15, 793–795.

Nogales, E., Scheres, S.H.W., 2015. Cryo-EM: A Unique Tool for the Visualization of Macromolecular Complexity. Mol. Cell 58, 677–689.

Paszke, A., Gross, S., Massa, F., Lerer, A., Bradbury, J., Chanan, G., Killeen, T., Lin, Z., Gimelshein, N., Antiga, L., Desmaison, A., Köpf, A., Yang, E., DeVito, Z., Raison, M., Tejani, A., Chilamkurthy, S., Steiner, B., Fang, L., Bai, J., Chintala, S., 2019. PyTorch: An Imperative Style, High-Performance Deep Learning Library. arXiv [cs.LG].

Peck, J.V., Fay, J.F., Strauss, J.D., 2022. High-speed high-resolution data collection on a 200 keV cryo-TEM. IUCrJ 9, 243–252.

Punjani, A., Rubinstein, J.L., Fleet, D.J., Brubaker, M.A., 2017. cryoSPARC: algorithms for rapid unsupervised cryo-EM structure determination. Nat. Methods 14, 290–296.

Rice, W.J., Cheng, A., Noble, A.J., Eng, E.T., Kim, L.Y., Carragher, B., Potter, C.S., 2018. Routine determination of ice thickness for cryo-EM grids. J. Struct. Biol. 204, 38–44.

Röder, C., Kupreichyk, T., Gremer, L., Schäfer, L.U., Pothula, K.R., Ravelli, R.B.G., Willbold, D., Hoyer, W., Schröder, G.F., 2020. Cryo-EM structure of islet amyloid polypeptide fibrils reveals similarities with amyloid-β fibrils. Nat. Struct. Mol. Biol. 27, 660–667.

Sanchez-Garcia, R., Segura, J., Maluenda, D., Sorzano, C.O.S., Carazo, J.M., 2020. MicrographCleaner: A python package for cryo-EM micrograph cleaning using deep learning. J. Struct. Biol. 210, 107498.

Tan, Y.Z., Aiyer, S., Mietzsch, M., Hull, J.A., McKenna, R., Grieger, J., Samulski, R.J., Baker, T.S., Agbandje-McKenna, M., Lyumkis, D., 2018. Sub-2 Å Ewald curvature corrected structure of an AAV2 capsid variant. Nat. Commun. 9, 3628.

Tegunov, D., Cramer, P., 2019. Real-time cryo-electron microscopy data preprocessing with Warp. Nat. Methods 16, 1146–1152.

Wagner, T., Merino, F., Stabrin, M., Moriya, T., Antoni, C., Apelbaum, A., Hagel, P., Sitsel, O., Raisch, T., Prumbaum, D., Quentin, D., Roderer, D., Tacke, S., Siebolds, B., Schubert, E., Shaikh, T.R., Lill, P., Gatsogiannis, C., Raunser, S., 2019. SPHIRE-crYOLO is a fast and accurate fully automated particle picker for cryo-EM. Commun Biol 2, 218.

Watkins, M.B., Wang, H., Burnim, A., Ando, N., 2023. Conformational switching and flexibility in cobalamin-dependent methionine synthase studied by small-angle X-ray scattering and cryoelectron microscopy. Proc. Natl. Acad. Sci. U. S. A. 120, e2302531120.

Wightman, R., 2019. PyTorch Image Models. GitHub repository. 10.5281/zenodo.4414861

Yadav, S.S., Jadhav, S.M., 2019. Deep convolutional neural network based medical image classification for disease diagnosis. Journal of Big Data 6, 1–18.

Yang, Y., Arseni, D., Zhang, W., Huang, M., Lövestam, S., Schweighauser, M., Kotecha, A., Murzin, A.G., Peak-Chew, S.Y., Macdonald, J., Lavenir, I., Garringer, H.J., Gelpi, E., Newell, K.L., Kovacs, G.G., Vidal, R., Ghetti, B., Ryskeldi-Falcon, B., Scheres, S.H.W., Goedert, M., 2022. Cryo-EM structures of amyloid-β 42 filaments from human brains. Science 375, 167–172.

Zheng, S.Q., Palovcak, E., Armache, J.-P., Verba, K.A., Cheng, Y., Agard, D.A., 2017. MotionCor2: anisotropic correction of beam-induced motion for improved cryo-electron microscopy. Nat. Methods 14, 331–332.

