## Supplementary Information for "Miffi: Improving the accuracy of CNN-based cryo-EM micrograph filtering with fine-tuning and Fourier space information"

Da Xu^1^, Nozomi Ando^1†^

^1^ Department of Chemistry and Chemical Biology, Cornell University, Ithaca, NY 14850, USA.

|  | Film | | | | | Drift | | | | | | Crystalline | | | | | | Contamination | | | | |
| --- | --- | --- | --- | --- | --- | --- | --- | --- | --- | --- | --- | --- | --- | --- | --- | --- | --- | --- | --- | --- | --- | --- |
|  | acc. | TP | FP | TN | FN | acc. | TP | FP | TN | FN | acc. | | TP | FP | TN | FN | acc. | | TP | FP | TN | FN |
| from scratch  single-channel | 0.876 | 3427 | 14 | 79 | 480 | 0.936 | 3279 | 214 | 463 | 44 | 0.927 | | 2916 | 288 | 792 | 4 | 0.999 | | 3994 | 3 | 2 | 1 |
| fine-tune  single-channel | 0.996 | 3893 | 0 | 93 | 14 | 0.988 | 3312 | 37 | 640 | 11 | 0.964 | | 2908 | 131 | 949 | 12 | 0.999 | | 3994 | 2 | 3 | 1 |
| fine-tune  two-channel | 0.999 | 3904 | 0 | 93 | 3 | 0.990 | 3307 | 25 | 652 | 16 | 0.993 | | 2909 | 18 | 1062 | 11 | 0.999 | | 3995 | 2 | 3 | 0 |

Table S1. Comparison of good-versus-bad prediction accuracy (acc.) within individual label categories of the validation set for the three models. True positive = TP; false positive = FP; true negative = TN; false negative = FN.

|  | Film | | | | | Drift | | | | | Crystalline | | | | | | Contamination | | | | |
| --- | --- | --- | --- | --- | --- | --- | --- | --- | --- | --- | --- | --- | --- | --- | --- | --- | --- | --- | --- | --- | --- |
|  | acc. | TP | FP | TN | FN | acc. | TP | FP | TN | FN | acc. | TP | FP | TN | FN | acc. | | TP | FP | TN | FN |
| EMPIAR-10175 | 0.987 | 965 | 1 | 42 | 12 | 0.962 | 862 | 32 | 119 | 7 | 0.997 | 1009 | 2 | 8 | 1 | 1.000 | | 1020 | 0 | 0 | 0 |
| EMPIAR-10344 | 0.990 | 2643 | 0 | 11 | 28 | 0.994 | 2545 | 10 | 122 | 5 | 0.990 | 2503 | 17 | 152 | 10 | 0.997 | | 2672 | 4 | 3 | 3 |
| EMPIAR-10379 | 0.996 | 1107 | 2 | 7 | 2 | 0.998 | 899 | 1 | 217 | 1 | 1.0 | 1113 | 0 | 5 | 0 | 0.996 | | 1112 | 1 | 2 | 3 |
| EMPIAR-10916 | 0.982 | 1709 | 0 | 22 | 31 | 0.998 | 1753 | 0 | 5 | 4 | 0.994 | 1752 | 0 | 0 | 10 | 0.996 | | 1754 | 0 | 0 | 8 |
| EMPIAR-11093 | 1.0 | 1330 | 0 | 0 | 0 | 0.992 | 1312 | 3 | 7 | 8 | 0.985 | 1144 | 17 | 166 | 3 | 0.969 | | 1289 | 0 | 0 | 41 |

Table S2. Good-versus-bad prediction accuracy (acc.) within individual label categories of additional test sets for the model trained with fine-tuning and two-channel input. True positive = TP; false positive = FP; true negative = TN; false negative = FN.

|  | acc. | TP | FP | TN | FN |
| --- | --- | --- | --- | --- | --- |
| EMPIAR-10175 | 0.747 | 673 | 113 | 89 | 145 |
| EMPIAR-10344 | 0.877 | 2160 | 113 | 191 | 218 |
| EMPIAR-10379 | 0.775 | 843 | 208 | 24 | 43 |
| EMPIAR-10916 | 0.849 | 1489 | 20 | 7 | 246 |
| EMPIAR-11093 | 0.926 | 1127 | 83 | 104 | 16 |

Table S3. Overall good-versus-bad accuracy (acc.) on additional test sets using MicAssess. True positive = TP; false positive = FP; true negative = TN; false negative = FN.

|  | cutoff | acc. | TP | FP | TN | FN |
| --- | --- | --- | --- | --- | --- | --- |
| EMPIAR-10175 | 8.03 Å | 0.821 | 788 | 153 | 49 | 30 |
| EMPIAR-10344 | 3.75 Å | 0.909 | 2368 | 233 | 71 | 10 |
| EMPIAR-10379 | 4.51 Å | 0.978 | 872 | 11 | 221 | 14 |
| EMPIAR-10916 | 4.53 Å | 0.951 | 1675 | 27 | 0 | 60 |
| EMPIAR-11093 | 6.68 Å | 0.840 | 1104 | 174 | 13 | 39 |

Table S4. Overall good-versus-bad accuracy (acc.) on additional test sets using CTF-based filtering. True positive = TP; false positive = FP; true negative = TN; false negative = FN.
